# A natural ANI gap that can define intra-species units of bacteriophages and other viruses

**DOI:** 10.1101/2024.04.18.590031

**Authors:** Borja Aldeguer-Riquelme, Roth E Conrad, Josefa Antón, Ramon Rossello-Mora, Konstantinos T. Konstantinidis

**Author notes:** Correspondence should be addressed to K.T.K.

## Abstract

Despite the importance of intra-species variants of viruses for causing disease and/or disrupting ecosystem functioning, there is no universally applicable standard to define these. A 95% whole-genome average nucleotide identity (ANI) gap is commonly used to define species, especially for bacteriophages, but whether a similar gap exists within species that can be used to define intra-species units has not been evaluated yet. Whole-genome comparisons among members of 1,016 bacteriophage species revealed a region of low frequency of pairs around 99.2-99.8% ANI, showing 3-fold or fewer pairs than expected for an even or normal distribution. This second gap is prevalent in viruses infecting various cultured or uncultured hosts, and from a variety of environments, although a few exceptions to this pattern were also observed (∼3.7% of the total species evaluated) and are likely attributed to cultivation biases. Similar results were observed for a limited set of eukaryotic viruses that are adequately sampled including SARS-CoV-2, whose ANI-based clusters matched well the WHO-defined Variants of Concern, indicating that they represent functionally and/or ecologically distinct units. The existence of sequence-discrete units appears to be predominantly driven by (high) ecological cohesiveness coupled to either recombination frequency for bacteriophages or selection and clonal evolution for other viruses such as SARS-CoV-2. These results indicate that fundamentally different underlying mechanisms could lead to similar diversity patterns. Based on these results, we propose the 99.5% ANI as a practical, standardized, and data-supported threshold for defining viral intra-species units of bacteriophages, for which we propose the term genomovars.

**Importance:** Viral species are composed of an ensemble of intra-species variants whose dynamic may have major implications for human and animal health and/or ecosystem functioning. However, the lack of universally-accepted standards to define these intra-species variants has led researchers to use different approaches for this task, creating inconsistent intra-species units across different viral families and confusion in communication. By comparing hundreds of viral bacteriophage genomes, we show that there is a nearly universal natural gap in whole-genome average nucleotide identities (ANI) among genomes at around 99.5%, which can be used to define intra-species units. Therefore, these results advance the molecular toolbox for tracking viral intra-species units and should facilitate future epidemiological and environmental studies.

## Introduction

Recognized viral species are often not homogeneous but consist of phenotypically and genotypically distinct variants (or intra-species populations), whose dynamics and evolution have major impacts on human and animal health, trophic webs, and ecosystem functioning. For example, the emergence of a new coronavirus variant in 2019 initiated the SARS-CoV-2 pandemic, which has killed an estimate of 6.95 million people across the world (World Health Organization, WHO). Beyond humans, the populations of European rabbits decreased by 60-70% when a new, more lethal variant of the rabbit hemorrhagic disease virus appeared (1). As a consequence, species that feed on rabbits, like the Iberian Lynx, followed the decline in rabbit populations in a clear example of how viral variants can influence trophic webs. Additionally, bacteriophages may exert population control of relevant environmental bacteria, such as *Synechococcus*, whose temporal genetic diversity co-varies with cyanophages due to viral infections (2). Additional studies have documented the constant rise and fall of genome variants as the underlying mechanism for the overall stable abundance of bacteriophage species over time (3, 4). Therefore, viral variants are apparently an important unit of viral diversity. However, a widely accepted definition of what a viral variant is and how much diversity it should encompass remain elusive.

The International Committee on Taxonomy of Viruses (ICTV) oversees the development, regulation, and maintenance of a universal taxonomic classification of viruses. However, the ICTV does not regulate the classification and nomenclature of organisms below the species level, nor does it provide a definition or criteria for intra-species units. The lack of a clear definition has led to confusion; consequently, the same (intra-species) terms, like strain or variant, have been frequently used with different standards and meanings. For instance, van Regenmortel defined strain as a virus with unique phenotypic characteristics (5). Accordingly, viruses with the same phenotype but different genomic sequences are considered to be the same strain, and the author reserves the term “variant” for these cases. Others, however, employed “variant” to distinguish between viruses with different phenotypes (6, 7), contrasting with the definition given by Van Regenmortel. The variety of definitions is also reflected in the diversity of criteria used to delineate intra-species units. Some authors used sequence identity values or clustering patterns of phylogenetic trees of single genes (8, 9) while others employed whole genome similarities, with a range of identity thresholds (98-100%)(10–12). It is important to note that the selection of the gene to use is typically an arbitrary decision, while different genes might produce different results (10). Furthermore, the whole genome sequence identity thresholds were primarily employed for practical reasons and convenience but their biological relevance in nature, if any, remains unknown.

Previous efforts have reported the existence of sequence-discrete viral units with intra-unit genome-wide ANI values being typically greater than 95%, a threshold that has been proposed as a standard for defining viral species (or vOTUs), especially bacteriophages (13–15). That is, the ANI values among genomes of the same species are higher than 95%, contrasting with <90% ANI to members of other species. Thus, there appears to be a natural gap in ANI values distribution between species. This is similar to the 95% ANI threshold commonly employed for microbial species definition (16, 17) and thus, sequence-discrete species seem to exist similarly for both microbes and their viruses. Recently, comparison of intra-species ANI values among genomes of the same bacterial species revealed the existence of a similar ANI gap between 99.2-99.8%, which has been proposed as a threshold to define intra-species units based on genomic data (18). Here, we aim to test whether a similar intra-species gap exists for viruses, which can be used to define or refine existing intra-species units, and assess the underlying molecular and/or ecological mechanisms for the any such gap.

## Results and Discussion

### An ANI gap within cultured viral species around 99.2-99.8%

We tested the existence of a similar ANI gap to that observed previously within prokaryotic species (18) among viral genomes using a dataset that included 75,012 prokaryotic virus (or bacteriophage) genomes and represented 306 viral species with a minimum of 20 genomes per species (Table S1). The ANI histogram based on all possible 51,522,103 intra-species pairwise genome comparisons, revealed a pronounced gap between 99.2-99.8% ANI (Figure 1A). To avoid the potential bias that could be introduced by a few well sampled species, we subsampled the data to the same number of pairs (150) per species, which corroborated the existence of the gap revealed based on the full data (Figure S1). For example, we found 854 values to fall within the 95.5% ANI bin vs 316 for the 99.5% ANI bin. We estimated that if the ANI values were randomly and evenly distributed between 95% and 100% ANI, we would have expected 847 comparisons per every 0.1% bin of ANI (43,200 pairs in total, divided by 51 bins between 95% and 100%). However, within the 99.4-99.6% bins there were only 374, 316 and 367 pairs, which is about 2.7 times fewer pairs than expected by chance. Indeed, bootstrapped (1,000) subsampling to the same number of pairs per species followed by automated peak and valley identification pointed to 99.3% as the deepest and most consistent across species valley in the ANI value distribution (Figure 1B-C).

**Figure 1.**
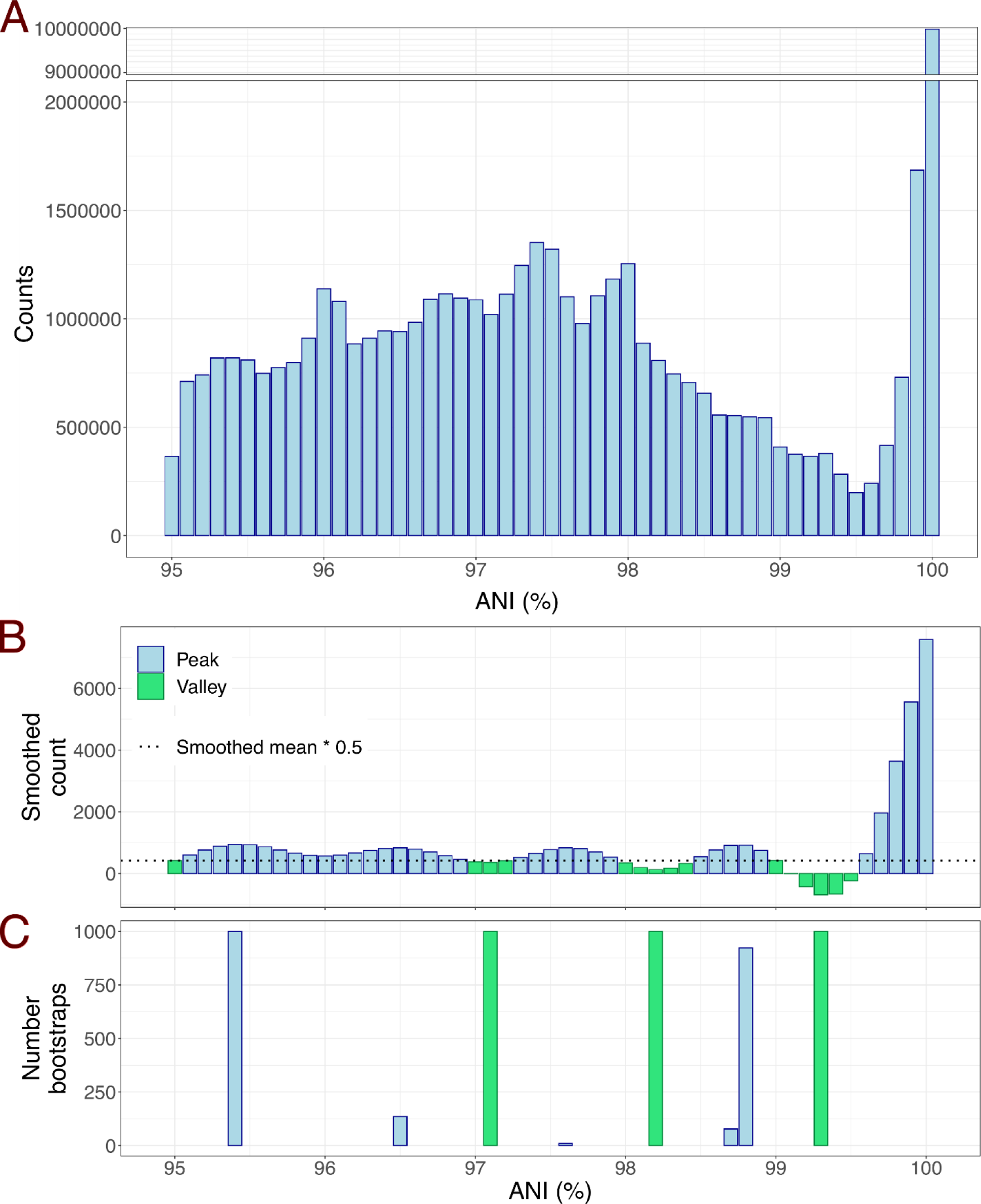
An intra-species ANI gap exists at around 99.2-99.8% and is consistently detected after 1,000 subsampling rounds. A) Histogram of ANI values without normalization to the same number of pairs per species. A total of 51,522,103 pairwise comparisons between 75,012 bacteriophage genomes were used. Note the pronounced drop in pairs between 99-99.8% ANI (∼300,000 per 0.1 units of ANI) compared to lower and higher ANI values (1,300,000 pairs at 97.4% ANI and 10,000,000 pairs at 100% ANI). B) Histogram of smoothed counts (sm_method=”gam”) using data subsampled to get the same number of pairs per species. Green bars indicate valleys while blue bars indicate intermediate values and peaks. C) Peaks (blue) and valleys (green) detected after 1,000 events of subsampling to 150 genome pairs per species and automatic peak and valley detection.

The species included in the dataset infect 609 different prokaryotic hosts species classified in 15 phyla, and be associated with animal or humans as well as environmental sources, which highlights the diversity of the dataset. Among the host species, *Escherichia coli* (24), *Mycobacterium smegmatis* (19), *Mycobacterium abscessus* (12) and *Serratia marcescens* (10) were the most predominant. Most of these viral species showed an intra-species ANI gap (see Figure S2 for examples, and https://github.com/baldeguer-riquelme/Viral-ANI-gap/ for all species evaluated). Indeed, 273 out of the 306 analyzed viral species (89.2%) showed an area of low frequency of pairs between 99.2-99.8% ANI (Groups 1 and 2). On the other hand, 26 species (8.5% of the total) did not show any peaks or valleys within the 99.2-99.8% ANI, which don’t provide evidence in favor or against the gap (undetermined distribution, Group 3) and only 7 species (2.5%) presented a contradictory distribution (Group 4: showing a peak rather than a valley around 99.2-99.8%) (Table 1, see Figure S3 for examples of each group). Therefore, these results revealed a nearly universal natural genomic threshold for distinguishing intra-species units.

**Table 1.** Classification of individual species based on the ANI distribution pattern. Group 1 refers to the species that show a valley between 99.2 - 99.8%, Group 2 includes species with predominately high-identity genomes or highly clonal (average ANI > 99.8%), Group 3 represents species with no peak or valley between 99.2 – 99.8% which is not consistent or contradictory with the gap (undetermined) and finally, Group 4 includes species showing a peak rather than a valley between 99.2– 99.8% (i.e., contradictory with the gap). The number of species within each group are shown, as well as the percentage they represent in parenthesis. *For long reads, we found the gap to be shifted towards lower ANI values (98.8 – 99.5%), so this range was employed to define the gap and an average ANI >= 99.5% was used to define species in Group 2.

|  | ANI gap |  | Group 3<br>(Undetermined) | Group 4<br>(Exceptions) | Total<br>species |
| --- | --- | --- | --- | --- | --- |
|  | Group 1<br>(Multiple<br>clusters) | Group 2<br>(1 cluster) |  |  |  |
| **Prokaryotic** | 231 (75.5%) | 42 (13.7%) | 26 (8.5%) | 7 (2.3%) | 306 |
| **Fosmid** | 7 (53.8%) | 2 (15.4%) | 4 (30.8%) | 0 (0%) | 13 |
| **Long reads\*** | 269 (38.6%) | 305 (43.8%) | 92 (13.2%) | 31 (4.4%) | 697 |

### Support for the ANI gap by culture-independent, long-read metagenomic data

To assess whether or not culture-independent data could provide further support to the existence of the ANI gap, we analyzed uncultured viral genomes recovered by fosmid sequencing and PacBio HiFi long-read metagenomes. Fosmids are cloning vectors of inserts up to 48 kbp long, and thus allow the recovery of complete or partial individual viral genomes. Similarly, PacBio HiFi provides high-quality consensus reads up to 25 kbp long. Therefore, both sequencing strategies could offer high resolution among co-occurring, uncultured viral genomes obtained directly from the environment with minimal sequencing error and bypassing isolation biases.

As shown in Figure 2, the analysis of the uncultured viral genomes recovered by fosmid or long-read sequencing also supported the existence of the ANI gap. Indeed, 9 out of the total 13 species adequately covered by fosmid sequencing presented the ANI gap, 4 species displayed an undetermined distribution, and none were inconsistent with the gap, similar to the results reported above for isolate genomes (Table 1). For 9 of these 13 species, we detected overlapping ends, indicating complete and circular genomes that truly represent single, distinct species rather than different regions of the same genome. The latter cannot be completely ruled out for the remaining 4 species. Fosmid metadata didn’t provide helpful information related to the taxonomy and/or ecology of the corresponding species, and thus we can only conclude that these species represented marine taxa/samples while their hosts remain unknown. Nevertheless, a previous study showed that these fosmids better represent the *in situ* abundant viruses than isolates (19) and are, most likely, a closer representation of the actual diversity in nature. Regarding long-read sequences, we did also observe a gap albeit slightly shifted towards lower ANI values (98.8-99.5%). Remarkably, 574 out of a total of 697 detected species in the long-read datasets displayed a clear ANI gap (Group 1: 269; Group 2: 305), 92 didn’t present a peak or valley at the gap (undetermined species), and only 31 showed an incompatible distribution (Table 1). The species displaying the ANI gap represented three distinct environments (i.e., human gut, chicken gut and seawater), which again supports its widespread existence in nature. Furthermore, long-read data mostly represent fragmented genomes, resulting in a higher dispersion of ANI and percentage of shared genome values around the mean, which presumably account for the slighted shifted range of the ANI gap mentioned above; nevertheless, the gap was evident even for these partial genomes (Figure 2B).

**Figure 2.**
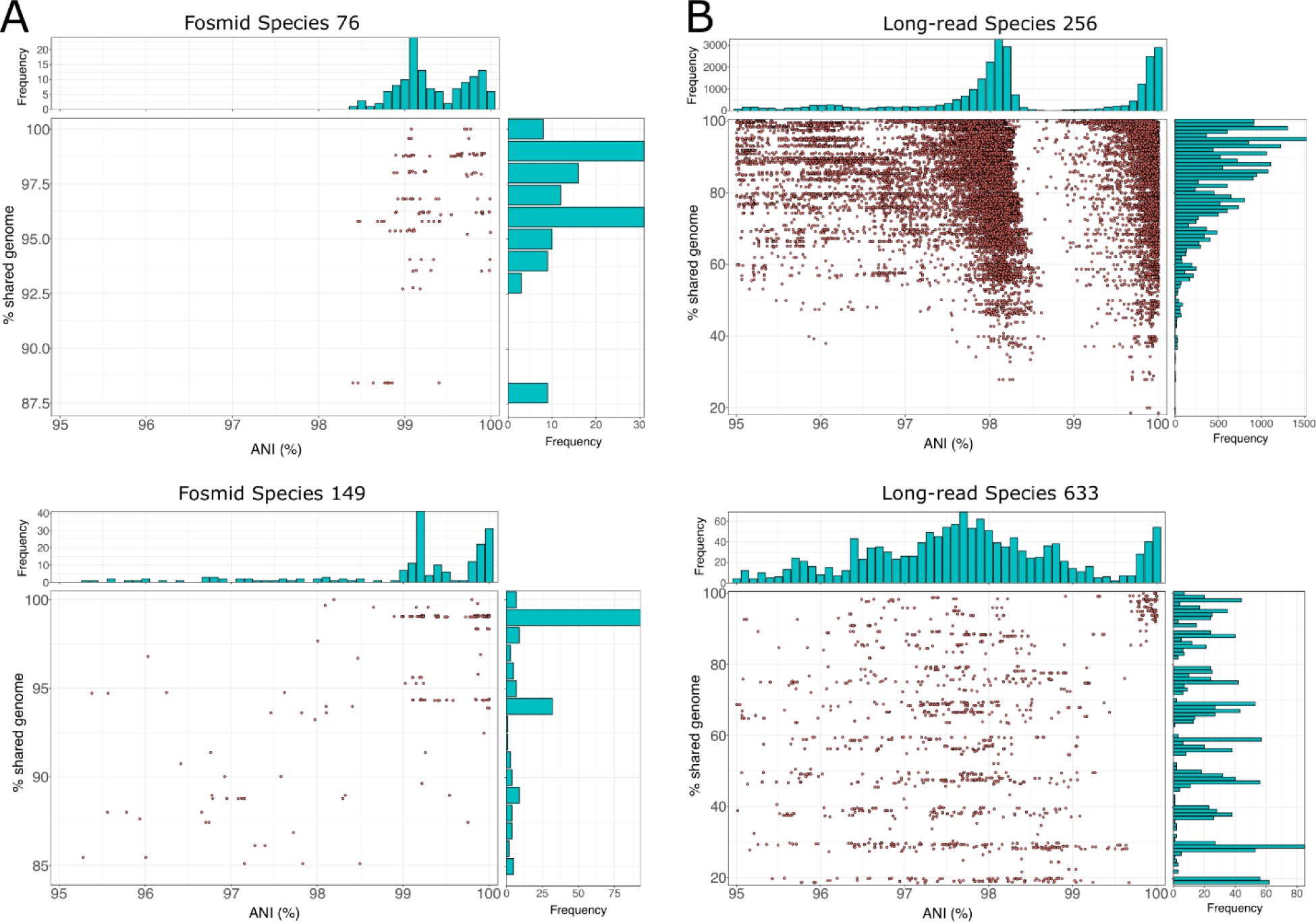
Percentage of shared genome (y-axis) vs ANI (x-axis) for two uncultured viral species recovered by fosmids (A) and long-read metagenomes (B). In the main panel of each figure, dots represent individual pairwise comparison between two genomes of the same species while histograms at the top and right side show the distribution of ANI values and the percentage of shared genome values, respectively, similar to Figure 1. Note the low frequency of pairs with ANI values around 99.5%. Plots for each viral species with more than 10 genomes can be found in https://github.com/baldeguer-riquelme/Viral-ANI-gap/.

### SARS-CoV-2 clusters defined by the ANI gap match WHO variants

The SARS-CoV-2 is probably the most sequenced virus to date, with more than 8 million genomes deposited in the NCBI database at the time of this writing. Furthermore, epidemiologic studies have provided detailed information on the phenotypic characteristics of the virus such as transmissibility or virulence of different viral variants. The WHO used these genotypic and phenotypic information to identify Variants of Concern (VOC), which include the most dangerous variants and the main drivers of the SARS-CoV-2 pandemic (20, 21). To date, five SARS-CoV-2 variants, named Alpha, Beta, Gamma, Delta and Omicron, have been declared as VOC by the WHO (22) and an additional one, named Epsilon, has been also declared by the Center for Disease Control and Prevention (CDC) of the United States (23).

To assess whether a similar ANI gap to the one revealed for bacteriophages above exists for human/animal viruses such as SARS-CoV-2, and minimize the potential bias introduced by different protocols, reagents, or assembly pipelines, we analyzed only the SARS-CoV-2 genomes deposited by the CDC (USA) and the Robert-Koch Institute (RKI, Germany), two of the main submitting institutions. Then, we randomly subsampled the dataset to get the same number of genomes for each one of the six VOC. The analysis of the CDC genomes showed a weak but evident signal of an ANI gap at around 99.8% (Figure 3A). When the analysis was restricted to high-quality genomes only (without any Ns), the gap was even more clearly observed (Figure 3B), and the ANI values were less dispersed especially at lower ANI values. This result highlights that undetermined positions (i.e., Ns) increase the noise of the ANI calculation, and thus high-quality genomes should be used for accurate results especially for relatively short genomes like SARS-CoV-2. The genomes sequenced by the RKI yielded similar results (Figure S4).

**Figure 3.**
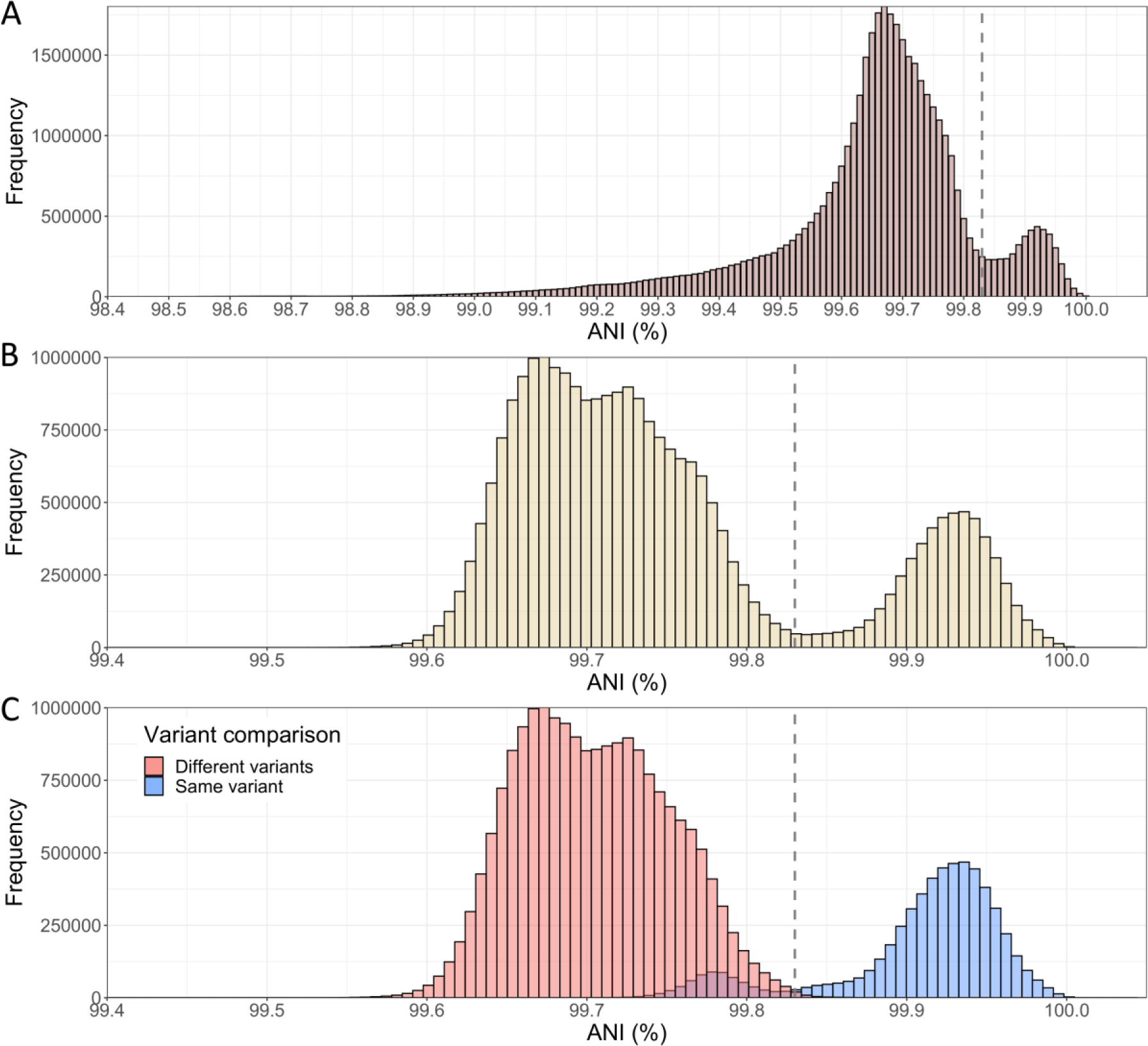
ANI histograms of the SARS-CoV-2 genomes sequenced by the US CDC. A) Histogram with a random subset of 6,481 genomes (Ns allowed) belonging to the Alpha (n=1250), Beta (n=534), Delta (n=1250), Gamma (n=1250), Epsilon (n=947) and Omicron (n=1250) VOCs were analyzed, that is 41,996,880 pairs in total. A subtle ANI gap at around 99.8% can be observed (see vertical line). B) Histogram with 5,041 high-quality genomes (i.e., without Ns) belonging to the Alpha (n=1250), Beta (n=99), Delta (n=1250), Gamma (n=1035), Epsilon (n=157) and Omicron (n=1250) VOCs were analyzed, that is 27,620,280 pairs in total. After removing genomes with Ns, the data reveal a clearer bimodal distribution and a more pronounced ANI gap for the SARS-CoV-2 genomes. C) Histogram shows the same data as in B, but the bars are colored based on whether the genomes compared are assigned to the same variants by WHO (in blue) or not (in red). Note the limited overlap between the two groups.

Remarkably, the distribution of genome pairs into the same or different variants overlapped, almost perfectly, with the highest and lowest peaks, respectively (Figure 3C). Considering the ANI value with the lowest number of pairs (99.83%) as a threshold to define variants, we found that 99.4% of all pairs above this threshold represented genomes belonging to the same variant. Conversely, only 3.2% of all pairs below 99.83% ANI involved genomes of the same variant. This result indicates that classification and identification of variants based on the intra-species ANI gap produces almost the same result as the combination of genotypic and phenotypic data employed by the international institutions. While the latter data are certainly needed for other purposes such as virulence assessment, the ANI gap offers a simple and complementary approach to identify viral variants. Furthermore, this result demonstrates that the resulting ANI-based clusters are associated with significantly distinct phenotypic characteristics. While we are able to confirm the latter only for the SARS-CoV-2 based on the abovementioned results, we hypothesize that clusters defined by the ANI gap within other viral species may also present significant phenotypic and/or ecological differences.

In addition to SARS-CoV-2, the existence of the ANI gap was also examined for a small set of eukaryotic viruses that have been adequately sampled. Similarly to the results reported above for bacteriophages, the gap was observed for a variety of viruses that infect humans, mosquitoes, birds, ruminants or pigs (Figure S5). Given the small size of the eukaryotic dataset and the fact that different thresholds for intra-species units are frequently used for these viruses compared to bacteriophages (24), the results reported here for the eukaryotic genome dataset can not be broadly extrapolated to all (or most) eukaryotic viruses. However, the fact that at least some eukaryotic viruses display the gap indicates that the intraspecies gap might be a widespread feature of viruses, not only bacteriophages. Interestingly, a threshold of 1% and 0.5% nucleotide differences between genomes was previously proposed to define human *Alphapapillomavirus* lineages and sublineages, respectively (25) and our 99.5% ANI threshold closely matches WHO’s designation of VOCs for SARS-Co-V2. Future research should more rigorously assess the existence of an intra-species ANI gap in eukaryotic viruses by sequencing -for instance-a broader range of species.

### Definition of intra-species clusters: a proposal for the term genomovar

Our results show that intra-species units of bacteriophages can be distinguished based on their ANI values. Given the multiple definitions of what constitutes a viral strain or a genetic variant, we propose the 99.5% ANI clusters to be referred to as genomovars. This term was proposed decades ago to distinguish bacterial genomic groups that belong to the same species but show distinct phenotypic and genomic features (26, 27). It has been recently proposed to refer to the intra-species clusters of prokaryotic organisms that share more than 99.5% ANI (28). Therefore, our proposal to use the same term for both viral and bacterial intra-species clusters will provide consistency and facilitate communication. We suggest the midpoint value (i.e., 99.5% ANI), rather than the upper value (i.e., 99.8% ANI) of the gap as a more conservative threshold and in order to account for the variation in ANI gap values observed among different species. However, this should only be considered as a practical and convenient standard to define intra-species units and genomovars, and researchers are encouraged to adjust this threshold to better match the ANI value distribution revealed by the data of their phage of interest.

### What are the underlying mechanism(s) for the 99.5% ANI gap?

Viral genome recombination may occur when two viruses co-infect the same cell, a phenomenon that has been observed in up to 50% of the total infected cells in the surface of the oceans and other environments (29). Once both genomes are inside the cell, recombination can take place coupled to replication (30), generating the widely described mosaicism of viral genomes (31). Since recombination is apparently an important mechanism that could drive viral genome evolution (30) we examined if it could be the underlying mechanism that maintains the intra-species ANI gap. Specifically, we tested the hypothesis that recent recombination is more frequent, and unbiased across the genome, within a cluster (e.g., a species or a genomovar) vs. between clusters of genomes, and thus can serve as the mechanism of cohesion for the cluster. For this, we first measured the fraction of genes displaying >99.8% identity between genome pairs (F100) relative to the expected number of such high-identity genes based on their ANI value and assuming no recombination (F100 expected; see Methods for details), as a proxy for recent recombination events (Figure 4A). For this analysis, we focused on the *Sal. ruber* phage species that more genomes are available from a natural population (32). The dataset included 177 high-quality viral genomes able to infect the same host (*Sal. ruber* strain M8) that were recovered from two adjacent ponds of the same saltern, sampled (just) two weeks apart in 2014. ANI distribution values highlighted a gap at 99.6%, consistent with the data reported above for all viral species (Figure 1), which translated to nine distinct 99.6% ANI-based genomovars. Genomes sharing <99.6% ANI shared more high-identity genes than expected based on the model with no recombination (Figure S6). Further, we calculated the fraction of recombinant genes between a reference genome and all genomes of different genomovars and observed that recombination occurred in 83 to 100% of the genes in the reference genome, depending on the genome used as reference (Figure 4B, Figure S7). Therefore, recombination is not only frequent enough, but also occurs throughout the entire genome.

**Figure 4.**
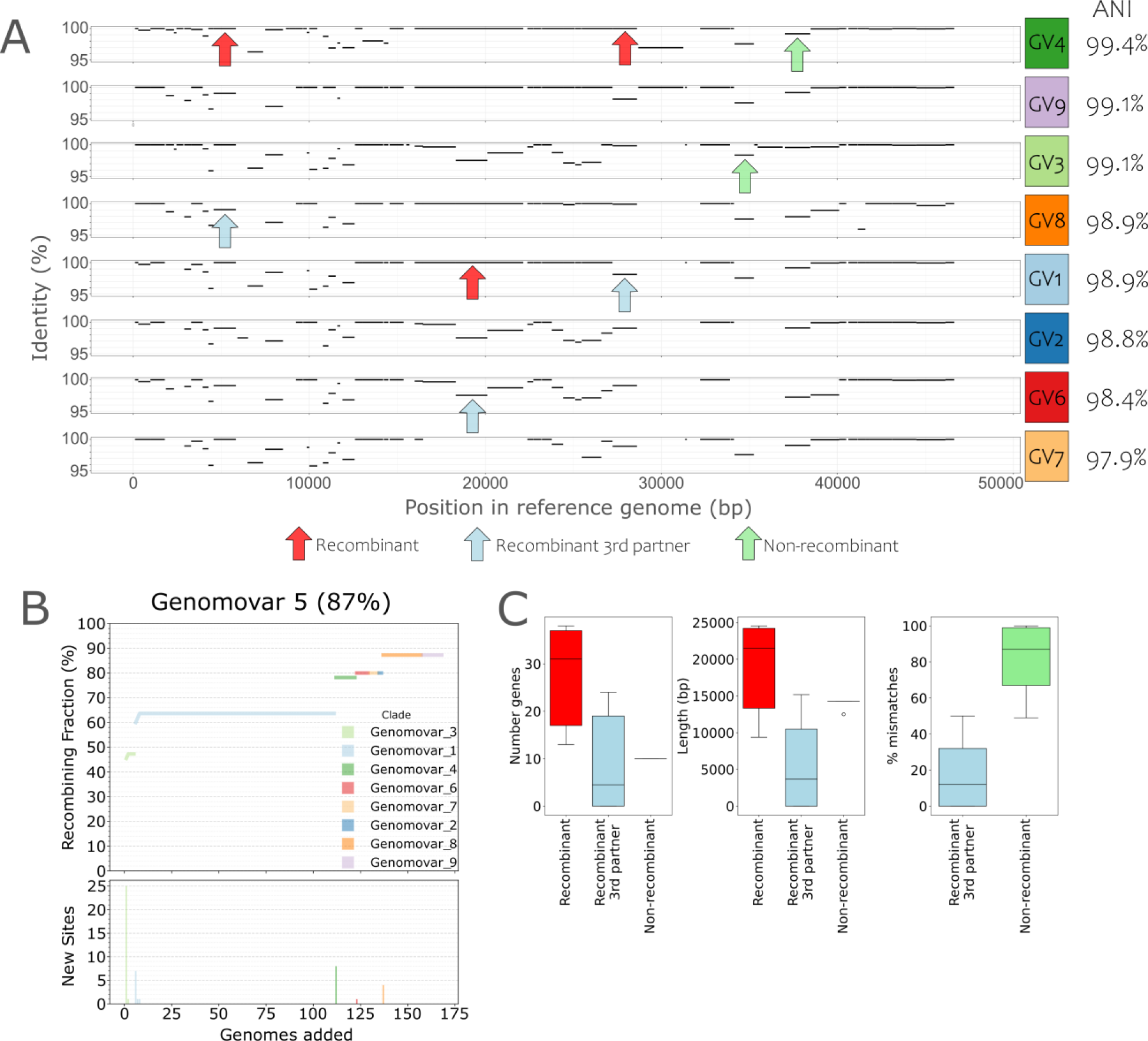
Quantifying the role of recombination as a force of cohesion of the intra-species units. A) Gene identity plot of a reference genome (genome 13v8 of genomovar 5) against query genomes of different genomovars. Plots are sorted by decreasing ANI against the reference genome (rightmost value). Each black line in the plot represents the position of a gene of the reference genome (x-axis) shared by the query genome and the percent identity of the shared homolog (y-axis); no lines represent genes that are specific to the reference genome (no match in the query genome). For each pair of genomes, shared genes with identities above 99.8% were considered as (recently) recombinant (red arrows, recombination as a force of cohesion); genes with identities below 99.8% within the pair of genomes in question but above 99.8% with another genome were classified as recombinant with a 3^rd^ partner (blue arrows, recombination as a force of diversification); genes with identities below 99.8% for all analyzed pairs were considered non-recombinant (green arrows, representing recent point mutations). B) Cumulative curve of the number of genes of the reference genome found to have recently recombined with a genome of another genomovar when adding the latter genomes sequentially in the analysis (x-axis). Top plot shows the number of recombined genes as a fraction of the total genes in the reference genome while the bottom plot shows the number of recombined genes newly detected by each genome added to the analysis. C) Quantification of the strength of recombination as a force of cohesion (red) and diversification (blue) following the gene classification described in panel A. Note that the impact of cohesive recombination, measured by the number of recombinant genes (left) or their total length (middle), is greater than diversifying recombination. GV: Genomovar.

Since a recombination event could increase similarity between the two partner genomes but, it could also increase dissimilarity between the two when recombination occurs with a third genome that is more divergent, recombination can be both a force of cohesion and diversification. To assess the relative importance of these two forces, we classified genes into three groups for each pairwise genome comparison: recombinant genes (>99.8% identity within the genome pair considered, proxy for cohesion force), recombinant genes with a 3^rd^ partner (<99.8% identity within the genome pair considered but >99.8% identity with a 3^rd^ genome, proxy for diversification force) and non-recombinant genes (<99.8% identity within the genome pair considered, proxy for point mutation force) (Figure 4A). We randomly selected one reference genome from each genomovar and calculated the total number, length and mismatches of genes classified in the three categories described above (Figure 4C, Figure S8). We observed that mismatches between a pair of genomes mainly involved ‘recombinant genes with a 3rd partner’ rather than ‘non-recombinant genes’ because most of the low identity alleles between the pair of genomes evaluated had a high identity (>99.8% identity) match with another genome, of a different genomovar, in our collection. These results suggested that recent point mutations have a relatively small impact on the evolution of these genomes relatively to recombination. Further, the number and length of ‘recombinant genes’ were higher than that of ‘recombinant genes with a 3^rd^ partner’ for 5 out of 9 genomovars, while only 3 genomovars displayed higher numbers for ‘recombinant genes with a 3^rd^ partner’ and 1 genomovar showed similar values for both categories. While our empirical approach, most likely, underestimates the frequency of recombinant genes with a 3^rd^ partner because our genome collections does not cover the total diversity of the natural population, cohesive recombination had similar contribution to diversifying recombination for at least a couple of the genome pairs evaluated when we added the genes found in mutation group to the 3^rd^ partner group. These results do indicate that recombination as a cohesion force might be, overall, frequent enough. It is important to note that it is not possible to perform this type of analysis for members of the same genomovar due to the high identity across the whole genome (i.e., there is no signal over the background level of sequence identity to detect recombination). Consistent with this assumption, F100 values among members of the same genomovar (>99.6% ANI) fall inside the confidence interval of the model that assumes no recombination (see purple dots in Figure S6). However, since recombination between genomovars is frequent, recombination within the same genomovar is certainly expected, and is likely even higher in frequency, given also that recombination generally increases with increasing sequence identity of the recombining partners (33). Therefore, recombination might also be the main force of cohesion and evolution of genomovars of DNA viruses, in addition to the force of cohesion at the species level.

In contrast to the *Sal. ruber* bacteriophages mentioned above, we observed low frequency of recombination for SARS-CoV-2 genomes (e.g., F100 values fall inside the confidence interval of the model that assumes no recombination for most SARS-CoV-2 genomes; Figure S9), consistent with recent literature (34–36), even though SARS-CoV-2 genomes also show a clear ANI gap similar to *Sal. ruber* bacteriophages (Fig. 3). Thus, it is likely that mutation rather than recombination is the main genetic mechanism driving SARS-CoV-2 genome diversification. This finding suggests that different mechanisms (i.e., cohesive recombination vs. diversifying point mutation) could have similar results on the intra-species diversity patterns of DNA and RNA viruses, that is, the existence of an ANI gap. Furthermore, the study of the SARS-CoV-2 dynamics has shown the emergence of new and more infectious SARS-CoV-2 genomovars that replace the existing genomovars, in a clear example of ecological competition (37, 38). These results indicate that competition between genomovars (i.e. selection) may also be an important underlying mechanism for the ANI gap in at least some viruses like SARS-CoV-2. Accordingly, intermediate genomes (i.e., their ANI values fall within the gap) might present less competitive phenotypes, on their way to extinction by natural selection, or genomovars that thrive under other conditions and/or hosts, and hence possibly not adequately sampled.

### Conclusions

While the dataset analyzed here certainly under-samples total viral species diversity, it includes genomes for 1,016 species, from 3 out of the 6 viral realms. Thus, we believe that the dataset is large enough to allow some initial views of universal patterns of intra-species diversity in bacteriophages. ANI comparisons between these genomes revealed a gap of values between 99.2-99.8% for most species, including viruses with different characteristics (e.g., host, environment and nucleic acid material). Further, this bimodal distribution observed for the SARS-CoV-2 matched the classification of variants proposed by WHO, highlighting the accuracy and robustness of the ANI gap to distinguish viral variants, and suggesting that genomovars defined by this gap can carry distinct phenotypic (e.g., different virulence and/or infectivity) and/or ecological properties. Therefore, we consider the data presented here as strong evidence for the widespread existence of functionally-distinct intra-species units of bacteriophages. We propose the 99.5% ANI as a practical, convenient, standardized and data-supported threshold to define these bacteriophage intra-species units, and the term genomovar to refer to them. Our data support that this threshold can be useful for most bacteriophages species, but we also recognize that viral genome diversity is vast, and therefore the threshold may need to be adjusted for specific viral species. Several viral species in our dataset did not show this major ANI pattern presumably due to sampling bias or their true diversity, and they should be studied in the future to better understand the mechanisms driving intra-species diversity. Future research will also need to assess whether a similar pattern widely exists for eukaryotic viruses. Here, we provide evidence that at least several of the eukaryotic viruses do display an intra-species gap, and thus the results reported here, primarily based on bacteriophage genomes, may be more broadly applicable to other viruses. Our results also indicated that the ANI gap may be the result of either recombination (*Sal. ruber* bacteriophages) or selection-driven diversifying mutation (SARS-CoV-2 genomes), confirming earlier hypotheses that gene exchange (biological species concept) or ecology (ecological species concept) can both explain the appearance and maintenance of species and intra-species units. It would be interesting to study additional species in the future to advance our understanding of the relative importance of these two processes and their interplay. We expect that the proposed ANI methodology and threshold advance the set of genomic tools to define and track the units of intra-species diversity, and thus will facilitate future epidemiological and environmental studies.

## Methods

The bacteriophage genomes analyzed here were retrieved from JGI IMG/VR database, selecting only those recovered from prokaryotic isolate genomes to avoid any effects (noise) from the assembly of metagenomic reads. This dataset was complemented with viral isolates from “The Actinobacteriophage Database” (https://phagesdb.org/)(39) and NCBI using the following search string: “Viruses[Organism] NOT cellular organisms[ORGN] NOT wgs[PROP] NOT gbdiv syn[prop] AND (srcdb_refseq[PROP] OR nuccore genome samespecies[Filter]” (Table S1). Since there are no tools to confidently estimate viral genome completeness for all types of viruses (i.e., available tools work well for certain groups of viruses, such as Caudovirales), only genomes longer than 20 kbp were analyzed with the aim of reducing the noise introduced by very incomplete genomes. In addition, 177 recently published *Salinibacter ruber* bacteriophage genomes (32) were also included in the dataset. As a result, the final dataset comprised a total of 75,012 genomes longer than 20 kbp (Table S1).

Uncultured genome sequences of fosmids and long-read metagenomes were downloaded from the NCBI Genome and SRA databases, respectively (Table S1). All fosmid sequences are of marine origin and have been described previously (40, 41). Presumably due to the large amount of DNA required to perform long-read sequencing, we did not find any viral long-read metagenomes in public databases at the time of this writing. Instead, we used 19 PacBio HiFi cellular metagenomes from human and chicken guts as well as seawater samples described previously (42–46). Sequences were quality filtered using filtlong v0.2.1 (https://github.com/rrwick/Filtlong) with a minimum read length of 10 kbp and a minimum window quality of 99. Samples with at least 5,000 surviving reads were first analyzed by VirSorter2 v2.2.4 (47) to identify viral sequences and then, with checkv v1.0.1 (48) to further refine these sequences as viral- or host-derived. Only those reads with at least one identified viral gene and more viral genes than host genes were retained for further analysis.

The 8.15 million SARS-CoV-2 genomes were downloaded from NCBI (accessed on August 2^nd^, 2023) and then, to retain only high-quality genomes, sequences with any undetermined position (Ns) identified by the FastA.filterN.pl (content=0) script of the enveomics collection (49) were discarded. For the CDC sequences, we subsampled them to 1,250 genomes per VOC (except Epsilon and Beta VOCs that had only 157 and 99 available genomes, respectively), that is 5,041 genomes were used in total. The RKI dataset was subsampled to 300 genomes per VOC (except Gamma and Epsilon VOCs with 112 and 2 genomes, respectively), which provided 1,314 genomes in total. In addition to SARS-CoV-2, eukaryotic viral genomes retrieved from NCBI using the search explained above were also analyzed (Table S1).

ANI values between viral genomes were calculated using FastANI v1.33 (16) “Many to Many” mode with a fragment length of 1 kbp to account for the shorter viral genomes (relative to the default 3 kbp for microbial genomes). Viral genomes were assigned to the same species when sharing more than 95% ANI, a previously proposed threshold (14) that we also corroborated within our dataset (Figure S10). Finally, self-matches were removed, and plots were drawn in R using ggplot2 v3.4.2 (50).

To challenge the robustness of the gap with the prokaryotic and eukaryotic cultured viruses, we performed a subsampling analysis to get the same number of pairwise comparisons per species (150). The data were then smoothed using the smooth_data function (sm_method=”gam”) from the gcplyr R package v1.9.0 (51) and peaks and valleys were automatically identified using findpeaks from the pracma R package v2.4.4 (https://cran.r-project.org/package=pracma). This process was repeated 1,000 times, the results were pooled and finally plotted using ggplot2 (we used the Bootstrap_analysis.R script available in https://github.com/baldeguer-riquelme/Viral-ANI-gap/).

To classify the observed ANI patterns, we defined four groups based on the detection of valleys and peaks using the approach outlined above, the average ANI value of the collection of genomes analyzed (of the specific species of interest) and the average smoothed counts. Briefly, the data (ANI values) were first smoothed, and peaks and valleys were automatically detected, as explained above. Then, areas of low frequency of pairs were defined as those ANI bins with a number of smoothed counts below the smoothed mean * 0.5. To validate a valley, it had to be detected by the findpeaks function and be in an area of low frequency of pairs. This approach ensure that validated valleys are at the bottom of the ANI distribution. Group 1 included species displaying a validated valley between 99.2–99.8% ANI, and an average ANI below 99.8%. Species with an average ANI above 99.8% were classified into Group 2 and represented highly clonal species. Both Group 1 and Group 2 species provide support to the existence of the gap since the area between 99.2-99.8% display a low number of pairs. Species on Group 2 and Group 1 are then composed by one or several clusters, respectively. Group 3 included species that didn’t show a peak or a valley between 99.2%-99.8% ANI, and thus, doesn’t provide strong evidence against - or in favor of-the existence of the gap (undetermined distributions). Finally, there was a group of species that showed a peak rather than a valley between 99.2–99.8% ANI, which is inconsistent with the existence of the gap. These species were classified in Group 4. Selected examples of these groups are shown on Figure S3; and all plots are available on https://github.com/baldeguer-riquelme/Viral-ANI-gap/. We manually reviewed all 1,016 species plots and moved 56 species (9 prokaryotic, 6 fosmids and 41 long-read) to a different group than the species was automatically assigned to using the methodology described above.

Frequency of recombination was calculated based on the fraction of shared identical reciprocal best match genes between genome pairs (F100) relative to those expected based on the ANI value of the genome and assuming no recombination, which can be considered a proxy for recent recombination events. For this, genes were first predicted using Prodigal v2.6.3 and then reciprocal best match genes for each pairwise comparison were identified using BLASTn (v2.14.0). We assumed that genes sharing more than 99.8% identity represent recently recombined genes and labeled them accordingly. Finally, we defined the F100 value as the fraction of recombinant genes from the total number of reciprocal best match genes for each pairwise comparison. To build the simulated model that only considers random mutations and no recombination, we employed the script Simulate_population_genomes.py, available at https://github.com/rotheconrad/Population-Genome-Simulator. To resemble the available *Salinibacter ruber* phages genomes as much as this was possible, the simulated population was created with the following parameters: -n 100 -g 70 -c 1 -cr 90 -mu 690 -sd 150. Cumulative recombinant genes curves were built using the 03g_Recombinant_group_analysis.py script of the F100_Prok_Recombination pipeline. For each pairwise comparison, genes were classified into three groups: recombinant genes (>99.8% identity, proxy for cohesion force), recombinant genes with a 3^rd^ partner (<99.8% identity but >100% identity with a 3^rd^ genome, proxy for diversification force) and non-recombinant genes (<99.8% identity, proxy for mutation force) as descried in the main text. Note that the genes are labeled separately for each pairwise comparison and thus, the same gene might be classified in different categories depending on the genome pair analyzed. For example, a gene of genome A can be classified as ‘recombinant’ with genome B and as ‘recombinant with a 3^rd^ partner’ when compared to genome C. In addition, note that non-recombinant genes might actually be recombinant with a partner not included in the dataset. Classification and plots were carried out using the 03g_Recombinant_group_extra_code.py script available at https://github.com/baldeguer-riquelme/Viral-ANI-gap/. We used the F100 approach to detect evidence of recent recombination of SARS-CoV-2 genomes. Specifically, we compared the SARS-CoV-2 F100 values against a model with no recombination (Simulate_population_genomes.py script, parameters: -n 10 -g 12 -c 1 -cr 90 -mu 1180 -sd 2200).

## Acknowledgments

This work has been supported, in part, by the US National Science Foundation (Award No 1759831 and 2129823) to KTK and the METACIRCLE project (PID2021-126114NB-C41) funded by the Spanish Ministry of Science and Innovation to JA.

## Data availability

Prokaryotic, eukaryotic and fosmid viral genomes have been retrieved from IMG/VR, NCBI and “The Actinobacteriophage Database” (https://phagesdb.org/) (see Table S1 for accession numbers). Long-read metagenomes were downloaded from SRA (see Table S1 for accession numbers). A more detailed description of the procedure and R scripts used in this study, as well as plots for each individual species and the ANI-pattern-group they were assigned to, can be found on https://github.com/baldeguer-riquelme/Viral-ANI-gap/. Further technical details and scripts for the recombination analyses are available at https://github.com/rotheconrad/F100_Prok_Recombination/.

