## Supplementary material for "A natural ANI gap that can define intra-species units of bacteriophages and other viruses"

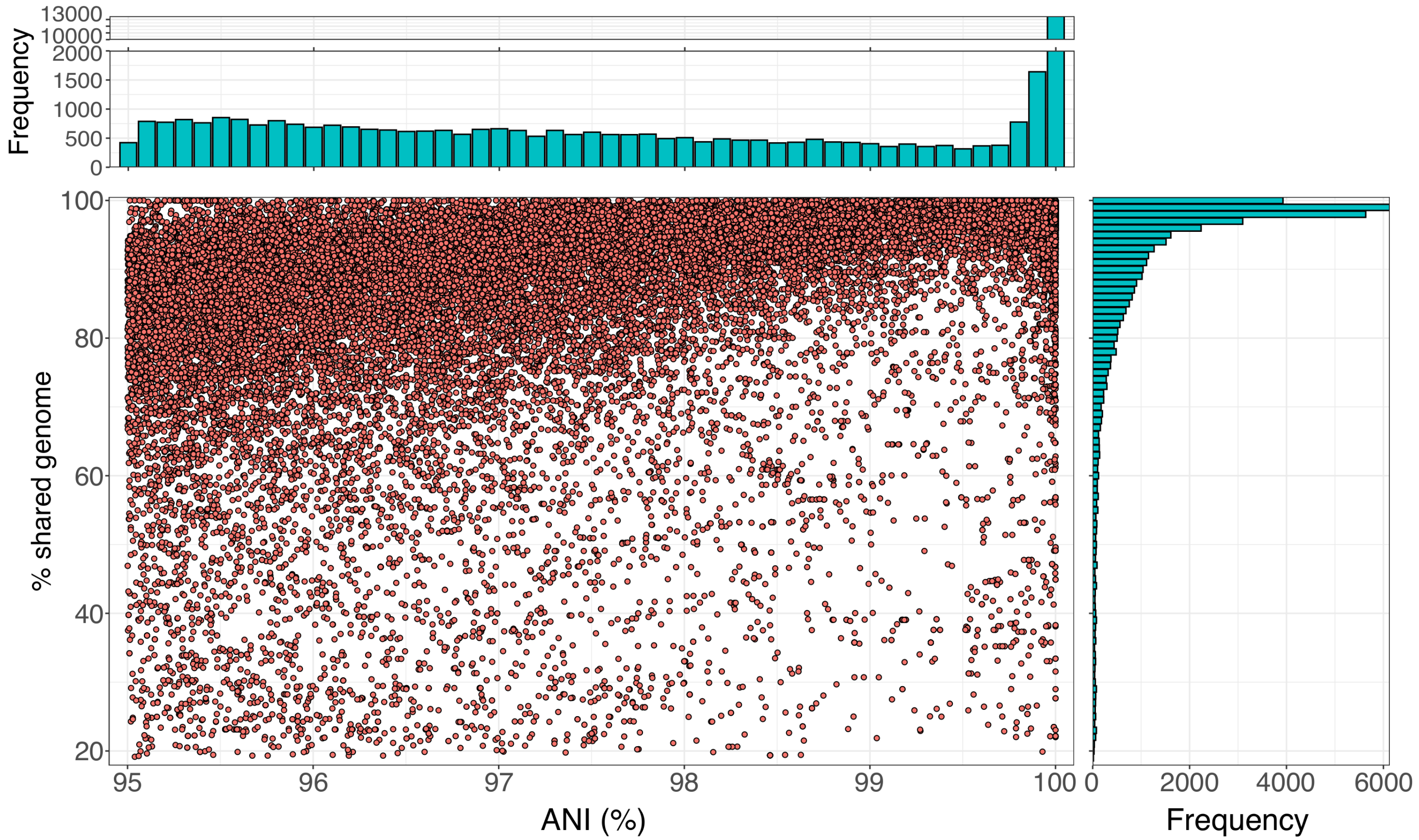


**Figure S1. Percentage of shared genome (y-axis) vs ANI (x-axis) for 288 bacteriophages species normalized to the same number of pairs per species.** In the main panel, dots represent individual pairwise comparisons between two genomes of the same species while histograms at the top and right side show the distribution of ANI values and the percentage of shared genome values, respectively. Only species with more than 20 genomes were included. To avoid biases due to the overrepresentation of a few species with many genomes, the underlying data was first subsampled to include the same number of pairs per species (n=150). Species with less than 150 pairs not included in the plot (18 species), this is why the number of species in this plot (288) differs from all species analyzed in the main text (306). Note the low frequency of pairs with ANI values around 99.2-99.8% relative to the number of values >99.8% or <99.2%.

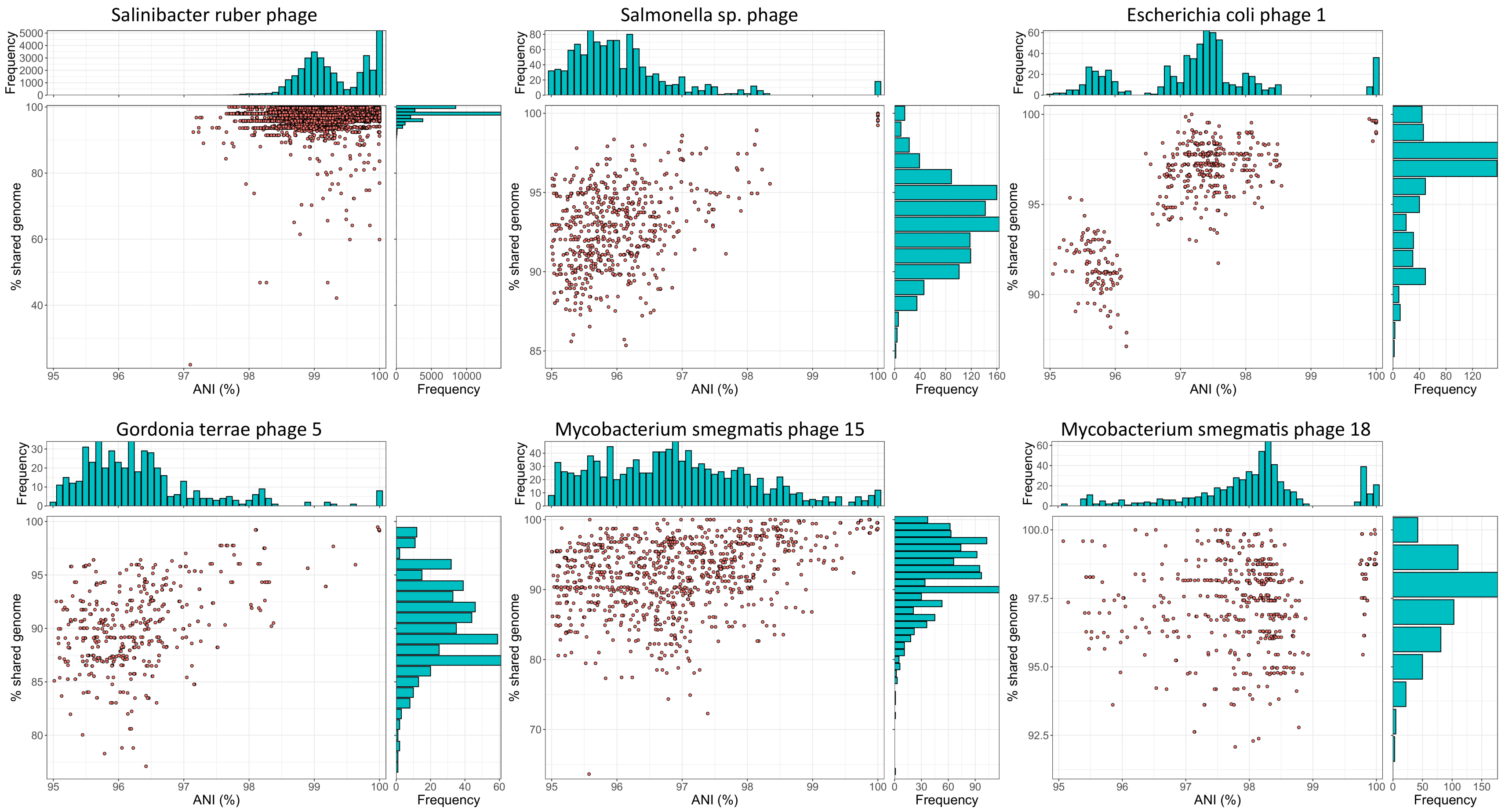


**Figure S2. Examples of individual viral species infecting prokaryotic hosts that show an ANI gap.** The plots are similar to those shown in Figures 1 and 2 but here data for individual species are shown instead. Plots for all individual species can be found at https://github.com/baldeguer-riquelme/Viral-ANI-gap/.

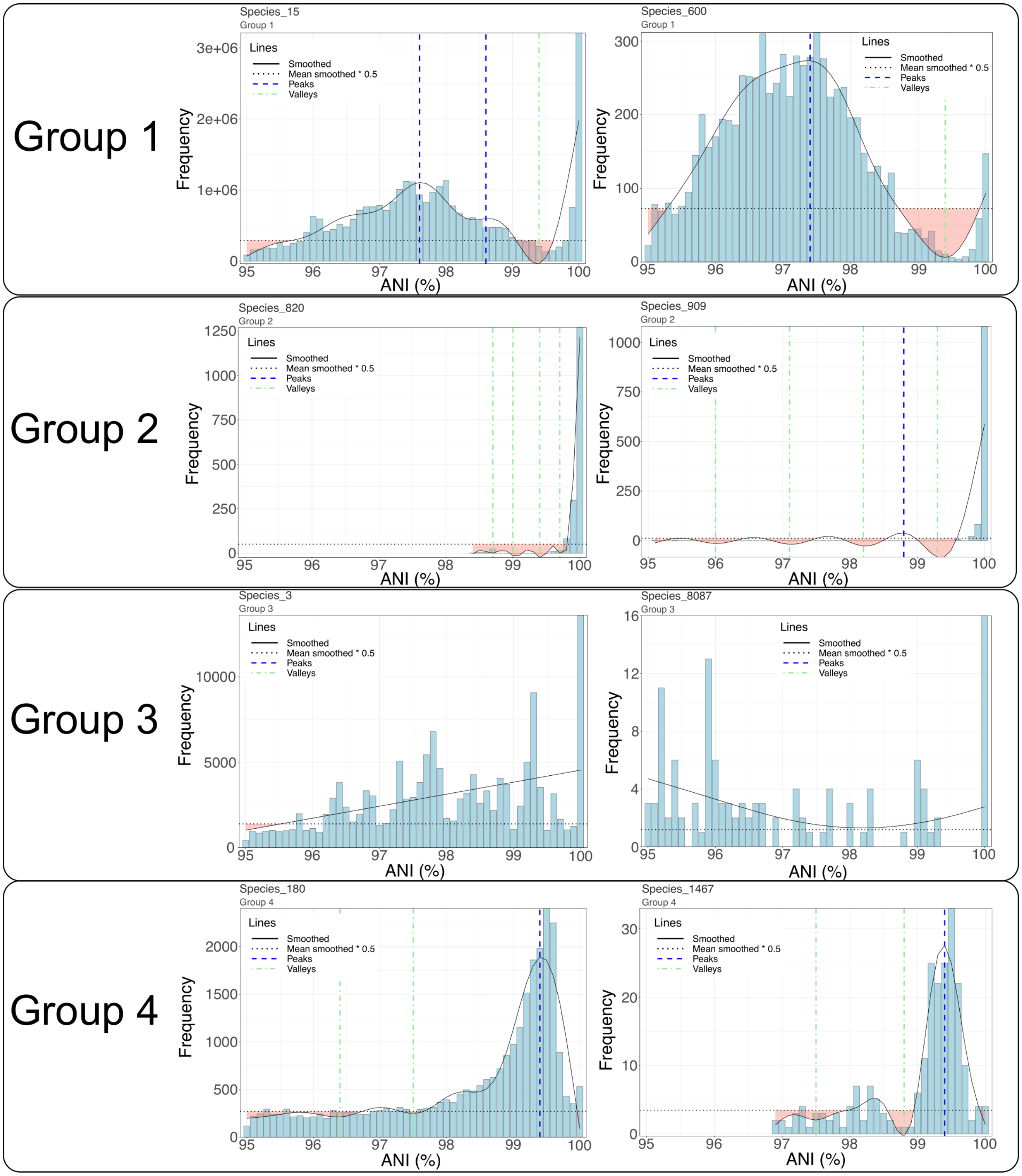


**Figure S3. Examples of the four groups used to classify viral species based on their intra-species ANI value distribution pattern.** Groups were defined based on the peaks and valleys detected, the average ANI value among the genomes of the group, and average smoothed counts. Group 1 included species displaying an ANI gap and were defined as those with a valley between 99.2–99.8% ANI, an average ANI below 99.8%. Species with an average ANI above 99.8% were classified into Group 2 and represented clonal species. Both Group 1 and Group 2, show areas of low frequency of pairs in the 99.2-99.8% range and the use of the ANI gap define several or one cluster for Group 1 and Group 2, respectively. Thus, these two groups provide evidence of the existence of a gap. Species in Group 3 didn't show a peak or valley between 99.2-99.8%. Thus, the ANI distribution of these species may reflect the actual, natural diversity pattern (not compatible with the gap) or a biased sequencing against highly similar genomes (compatible with the gap). We considered this species as undetermined. Finally, Group 4 included species that showed a peak rather than a valley between 99.2–99.8% ANI, which is not compatible with the existence of the gap.

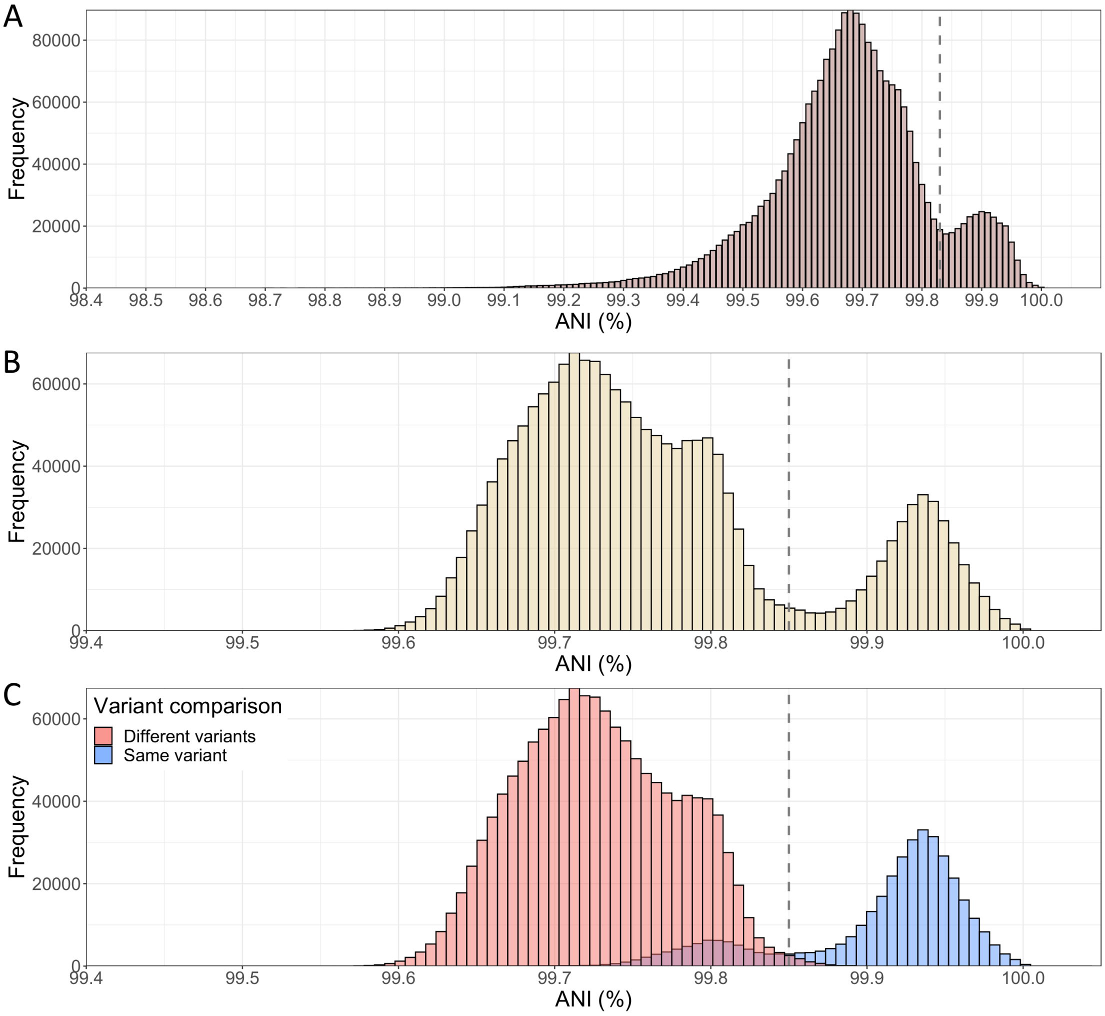


**Figure S4. ANI histograms of the SARS-CoV-2 genomes sequenced by the Robert-Koch Institute (Germany).** A) Histogram with a random subset of 1,506 genomes (Ns allowed) belonging to the Alpha (n=300), Beta (n=300), Delta (n=300), Gamma (n=300), Epsilon (n=6) and Omicron (n=300) VOCs were analyzed, that is 2,266,530 pairs in total. A subtle ANI gap at around 99.8% can be observed (see vertical line). B) Histogram with 1,314 high-quality genomes (i.e., without Ns) belonging to the Alpha (n=300), Beta (n=300), Delta (n=300), Gamma (n=112), Epsilon (n=2) and Omicron (n=300) VOCs were analyzed, that is 1,725,282 pairs in total. After removing genomes with Ns, the data reveal a clearer bimodal distribution and a more pronounced ANI gap for the SARS-CoV-2 genomes. C) Histogram shows the same data as in B, but the bars are colored based on whether the genomes compared are assigned to the same variants by WHO (in blue) or not (in red). Note the limited overlap between the two groups.

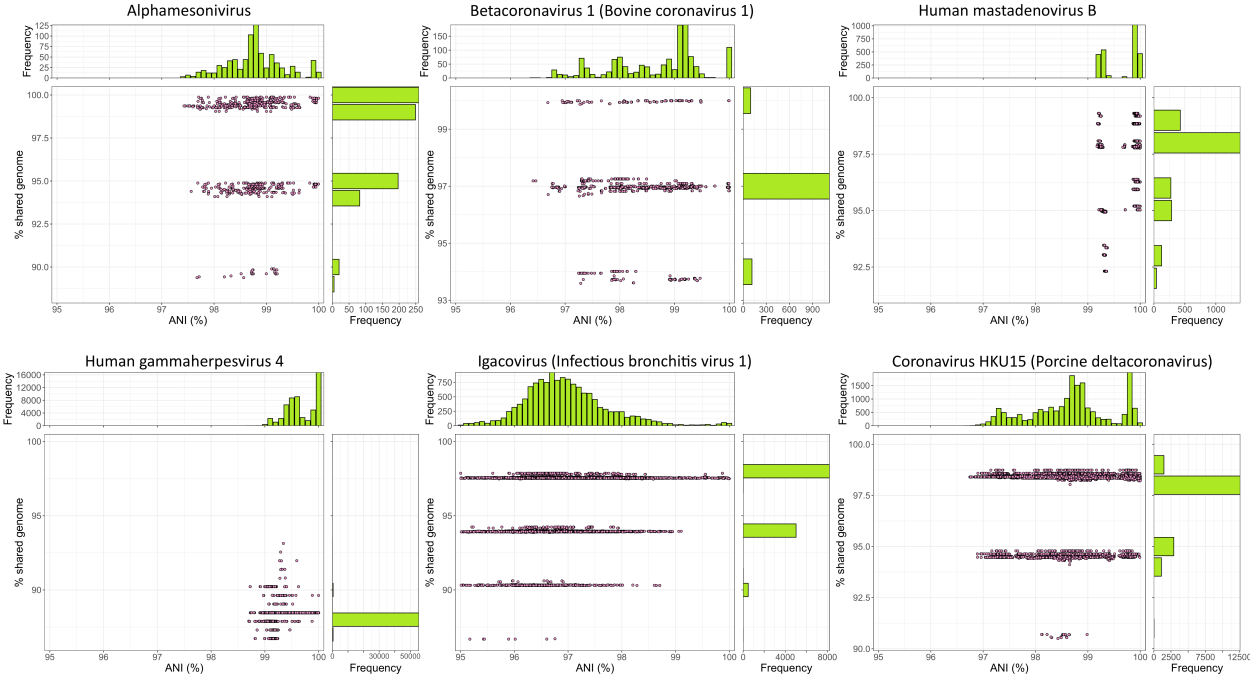


**Figure S5. Examples of individual viral species infecting eukaryotic hosts that show an ANI gap.** The plots are similar to those shown in Figure 1 and 2 but here data for individual species are shown instead. Note that the ANI gap is present for species with both RNA (Alphamesonivirus, Betacoronavirus 1, Igacovirus, Coronavirus HKU15) and DNA genomes (Human mastadenovirus B, Human gammaherpesvirus 4). Individual plots for all species can be found at https://github.com/baldeguer-riquelme/Viral-ANI-gap/.

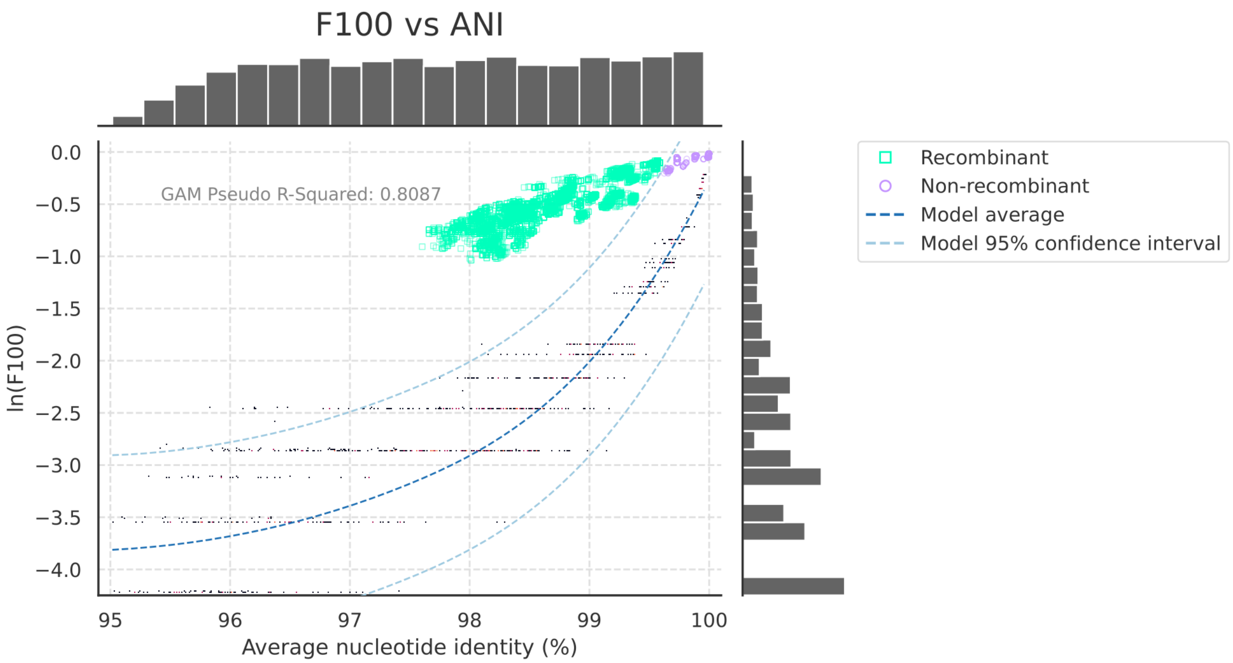


**Figure S6. Fraction of identical (>99.8% identity) reciprocal best match genes (F100, y-axis) as a function of the ANI (x-axis) of the genome pairs compared for *Salinibacter ruber* bacteriophages.** The fraction of identical genes (99.8% nucleotide sequence identity, F100 metric) and ANI values between two genomes compared against the number of such high identity gene expected under no recombination can be considered as a proxy for recent recombination events. Each dot or square represents the F100 and ANI values for a pair of genomes. Small black dots show the values for a simulated model with random mutation as the only evolutionary force (that is, simulated genomes with no recombination events between them). The data was fitted to a Generalized Additive Model (GAM) and the R-squared value is shown in the main panel; dashed lines show the mean (dark blue) and the 95% confidence interval (light blue) of the model. Purple dots and green squares represent the real data and indicate pairs with no more recombination than expected by the model and with more recombination than expected, respectively. The histograms at the top and right side of the plot show the number of ANI values and ln(F100) values for each bin unit, respectively.  Note that genome pairs sharing >99.5% ANI (same genomovar) show F100 values inside the confidence interval of the model, indicating that recombination events between such genomes cannot be distinguished from high sequence conservation.

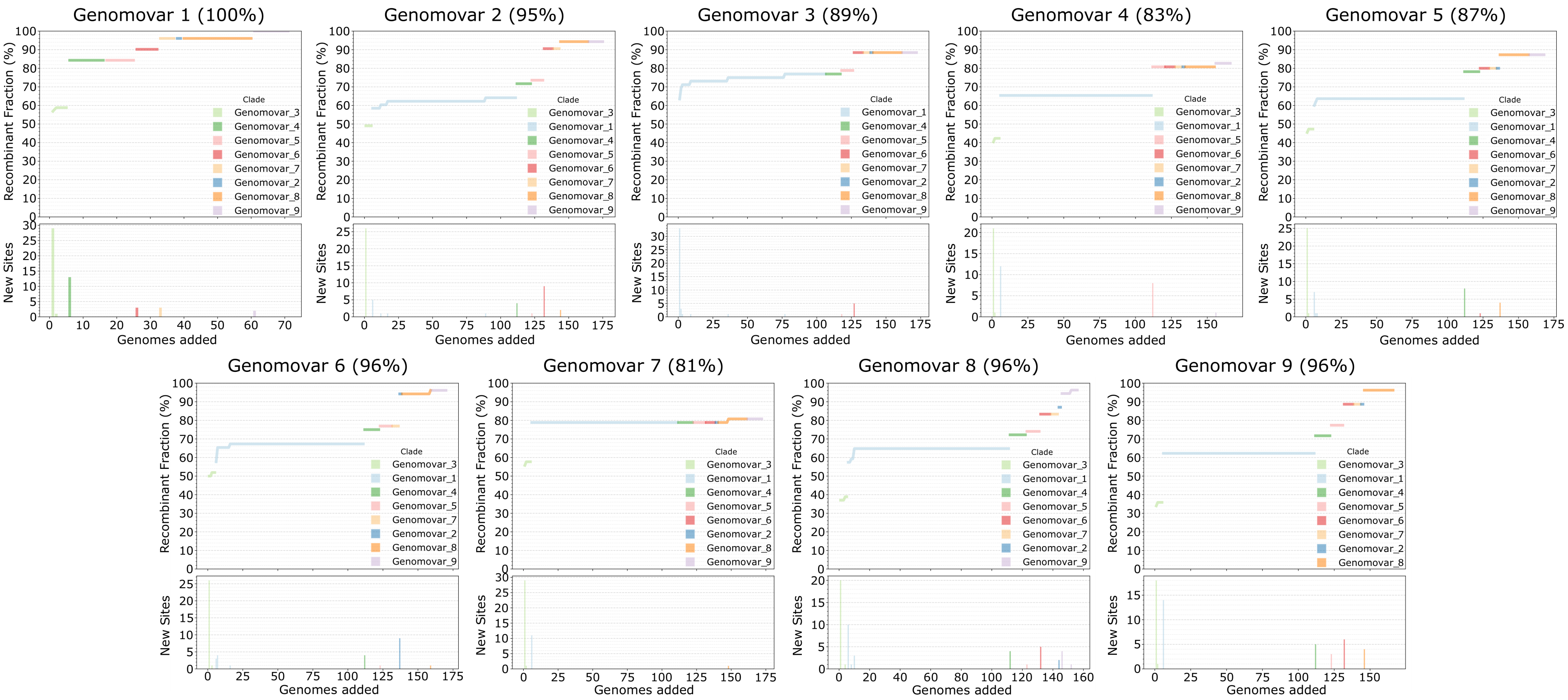


**Figure S7. Cumulative curve of recombinant genes.** Each panel shows the cumulative curve of the number of genes of one reference genome (graph title) found to have recently recombined with a genome of another genomovar (y-axis) when adding the latter genomes sequentially in the analysis (x-axis). Top plot shows the fraction of recombinant genes with respect to the total number of genes in the reference genome while the bottom plot shows the number of recombinant genes newly detected by each genome added to the analysis.

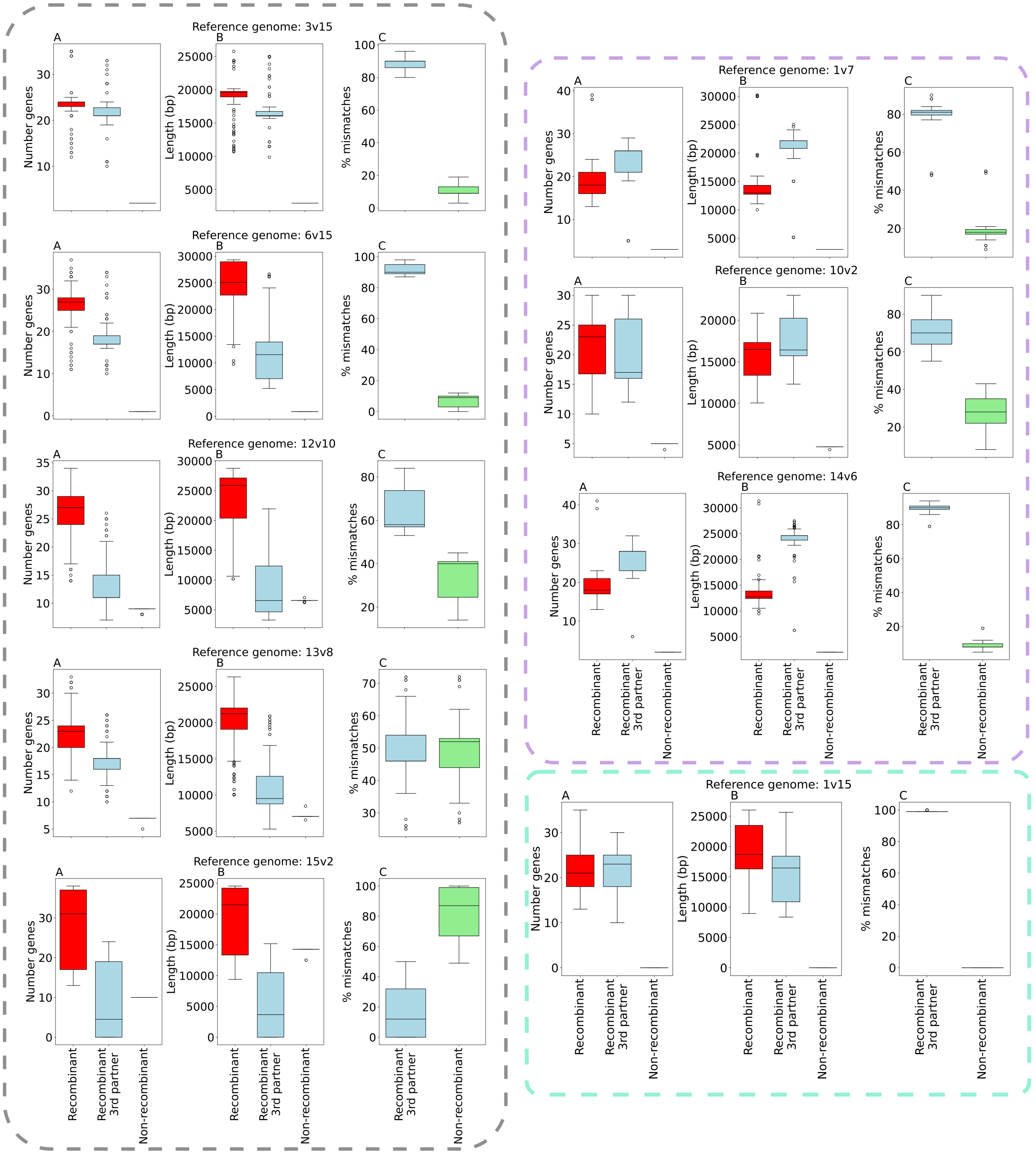


**Figure S8. Assessing the relative impact of recombination as a cohesive vs diversifying force.** Each row shows the results for pairwise comparisons of a reference genome of one genomovar against all genomes of different genomovars and consists of 3 plots that show the number of genes (A), their length (B) and the percentage of mismatches between each genome pair (C). For each plot, the genes were grouped into three categories: recombinant genes (red, proxy for cohesion force), recombinant genes with a 3rd partner (blue, proxy for diversification force) and non-recombinant genes (green, proxy for mutations). At the left, within a red rectangle with dashed lines, five genomovars that showed a higher strength of recombination as a force of cohesion are shown (red). At the top right, within a blue rectangle, three genomovars that showed higher strength of recombination as a force of diversification are shown. At the bottom right, within a green rectangle, one genomovar that showed equal strength of recombination as force of cohesion and diversification is shown.

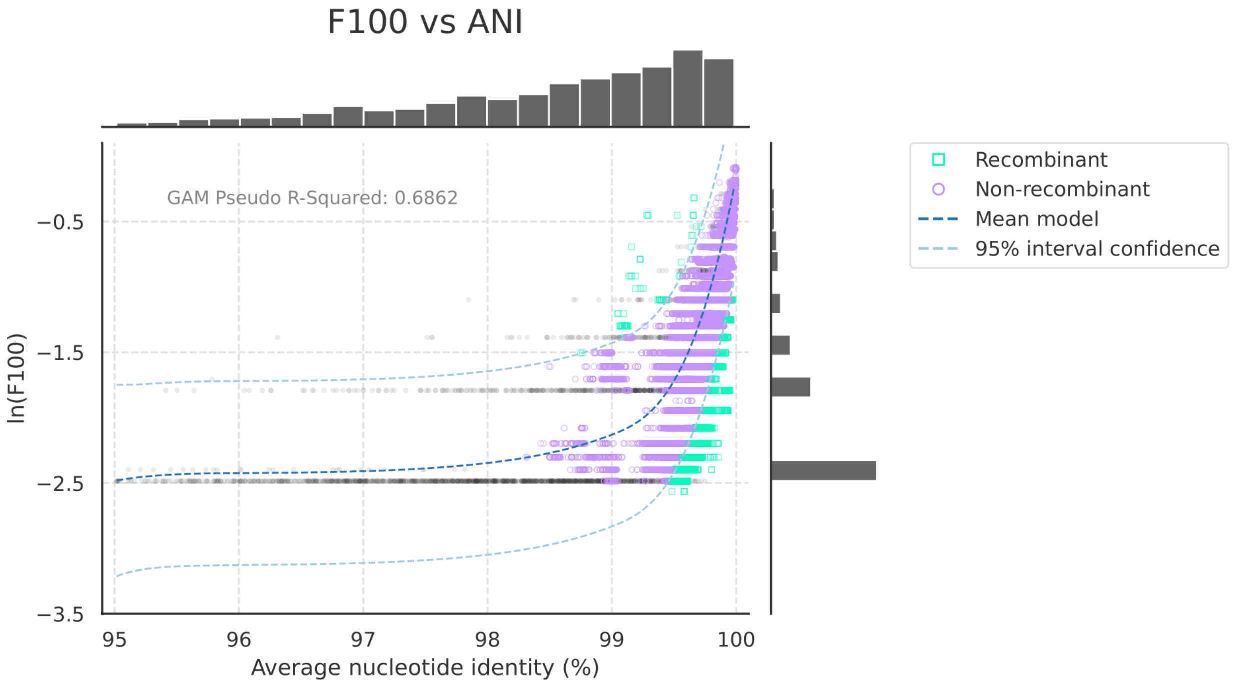


**Figure S9. Fraction of identical (>99.8% identity) reciprocal best match genes (F100, y-axis) as a function of the ANI (x-axis) of the genome pairs compared for SARS-CoV-2 genomes.** The fraction of identical genes (99.8% nucleotide sequence identity, F100 metric) and ANI values between two genomes compared against the number of such high identity gene expected under no recombination can be considered as a proxy for recent recombination events. Each dot or square represents the F100 and ANI values for a pair of genomes. Small black dots show the values for a simulated model with random mutation as the only evolutionary force (that is, simulated genomes with no recombination events between them). The data was fitted to a Generalized Additive Model (GAM) and the R-squared value is shown in the main panel; dashed lines show the mean (dark blue) and the 95% confidence interval (light blue) of the model. Purple dots and green squares represent the real data and indicate pairs with no more recombination than expected by the model and with more recombination than expected, respectively. The histograms at the top and right side of the plot show the number of ANI values and ln(F100) values for each bin unit, respectively. Note that for most genome pairs (99.9%) F100 values fall inside the confidence interval of the no recombination model, suggesting that no recent recombination events have occurred between those genomes. Only 0.07% of all pairs may be recombinant.

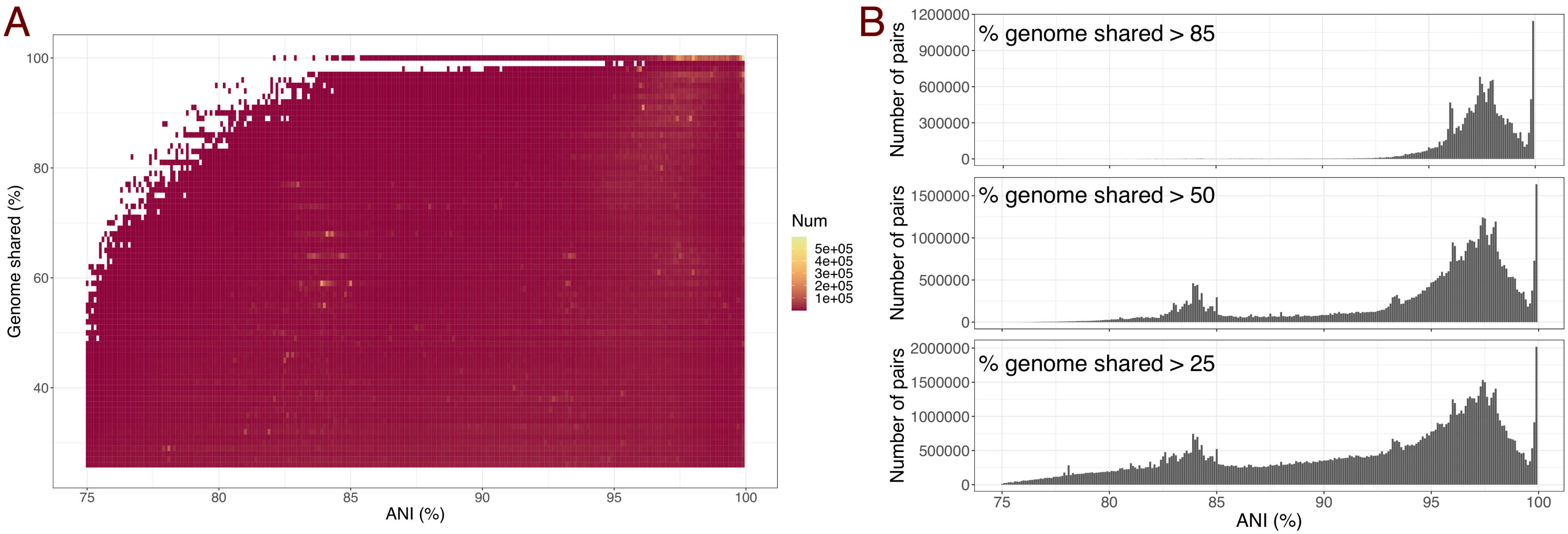


**Figure S10. An inter-species ANI gap exists between 85-95% ANI.** A) Heatmap of percentage of genome shared (y-axis) vs ANI values (x-axis). A total of 111,395,332 pairwise comparisons sharing more than 25% genome length were included in the plot. Note the existence of one cluster at >95% ANI and >80% percentage genome shared and another one at 82-85% ANI and 50-70% genome shared. B) Histograms of ANI values between 75-100% ANI for pairs sharing more than 85, 50 and 25% genome length. Note that the gap between 85-95% ANI is consistent for all three filters of minimum percentage of genome shared (>25, 50 and 85%). Indeed, pairs sharing almost their full genome length (>85%) only present identities above 95% ANI.
